# Optical Control of Microtubule Accumulation and Dispersion by Tau-Derived Peptide-Fused Photo-Responsive Protein

**DOI:** 10.1101/2024.09.24.614838

**Authors:** Soei Watari, Hiroshi Inaba, Qianru H Lv, Muneyoshi Ichikawa, Takashi Iwasaki, Bingxun Wang, Hisashi Tadakuma, Akira Kakugo, Kazunori Matsuura

**Affiliations:** Department of Chemistry and Biotechnology, Graduate School of Engineering, Tottori University, Tottori 680-8552, Japan; Center for Research on Green Sustainable Chemistry, Tottori University, Tottori 680-8552, Japan; State Key Laboratory of Genetic Engineering, Department of Biochemistry and Biophysics, School of Life Sciences, Fudan University, Shanghai, 200438, China; Department of Bioresources Science, Graduate School of Agricultural Sciences, Tottori University, Tottori 680-8553, Japan; School of Life Science and Technology, ShanghaiTech University, Shanghai 201210, China; Department of Physics and Astronomy, Graduate School of Science, Kyoto University, Kyoto 606-8502, Japan

## Abstract

Microtubules, a major component of the cytoskeleton consisting of tubulin dimers, are involved in various cellular functions, including forming axons and dendrites of neurons and retaining cell shapes by forming various accumulated superstructures such as bundles and doublets. Moreover, microtubule-accumulated structures like swarming microtubule assemblies are attractive components for dynamic materials, such as active matter and molecular robots. Thus, dynamic control of microtubule superstructures is an important topic. However, implementing stimulus-dependent control of superstructures remains challenging. This challenge can be resolved by developing designer protein approaches. We have previously developed a Tau-derived peptide (TP), which binds to the inner or outer surface of microtubules depending on the timing of the incubation. In this report, we designed the TP-fused photo-switchable protein Dronpa (TP-Dronpa) that reversibly photoconverts between monomeric and tetrameric states to photocontrol microtubule assemblies. The formation of microtubule superstructures, including bundles and doublets, was induced by tetrameric TP-Dronpa, whereas monomeric TP-Dronpa ensured that microtubules remained dispersed. Tetrameric TP-Dronpa also induced motile aster-like structures and swarming movement of microtubules on a kinesin-coated substrate. The formation/dissociation of these microtubule superstructures can be controlled by light irradiation. This system can generate and photocontrol various microtubule superstructures and provides an approach to facilitate the assembly of dynamic materials for various applications.

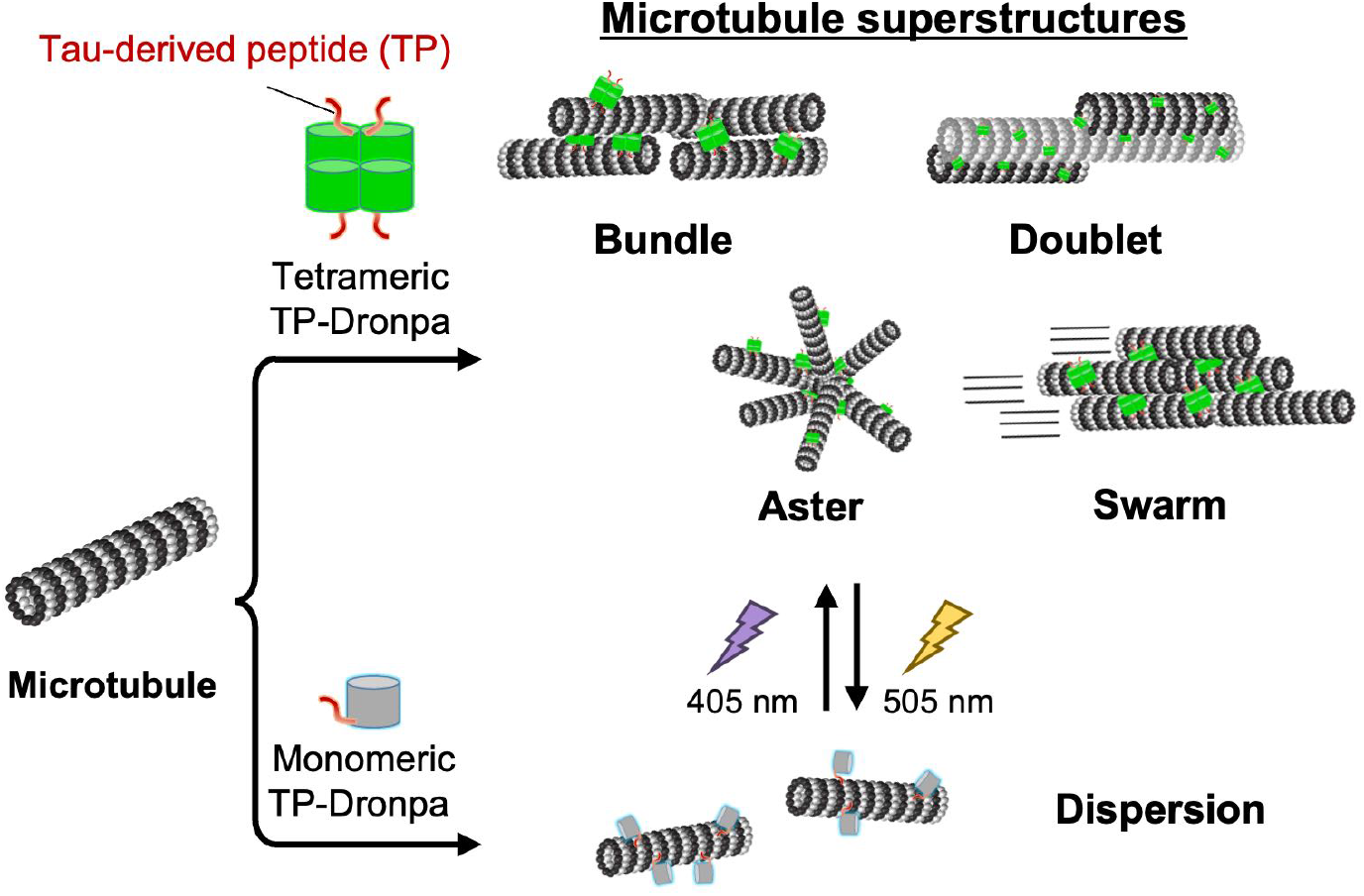

## INTRODUCTION

Microtubules are tubular cytoskeletons with micrometer lengths and a 15 nm inner diameter that form by polymerizing tubulin dimers with GTP and are involved in various functions, including cell division, cell shape regulation and cell migration.^1–4^ In nature, microtubule superstructures are formed by microtubules assembling to form bundles, doublet and aster-like structures. In neurons, microtubule-associated proteins (MAPs), such as Tau and MAP2, bundle microtubules by crosslinking microtubules to contribute to forming axons and dendrites.^5^ Microtubules also form doublet structures, which connect a complete microtubule to an incomplete microtubule in flagella and cilia to tolerate physical stresses such as waiving motion.^6–8^ Microtubules and associated superstructures play crucial roles as scaffolds of motor proteins such as kinesin and dynein for cargo delivery and the beating of flagella and cilia.^9^ The motile properties of microtubules are attracting attention for the construction of active matter and molecular robots.^10–13^ In particular, control of microtubule superstructures by external stimuli should facilitate the development of intelligent, active matter.^14^ For example, Kakugo et al. achieved optical control of a microtubule swarming motion by conjugating photochromic azobenzene-conjugated complementary DNA to the outer surface of microtubules.^15–17^ Moreover, Dogic et al. succeeded in the optical control of forming active gels consisting of microtubules and kinesin-fused iLID, a photo-responsive protein.^18^ In nature, regulating axon, dendrite and nucleus formation is conducted by various MAPs that crosslink microtubules.^5^ Mimicking the natural system would be useful for controlling microtubule superstructures. However, artificially designed proteins have not been reported for this purpose.

We have previously designed a Tau-derived peptide (TP: CGGGKKHVPGGGSVQIVYKPVDL) based on the repeat domain of Tau, which binds to the inner pocket of microtubules.^19^ TP binds to the inner surface of microtubules by preincubation with tubulin and subsequent microtubule polymerization, whereas TP binds to the outer surface of microtubules by incubation with polymerized microtubules.^19,20^ Conjugation of photoreactive molecules, diazirine^21^ and spiropyran/merocianine^22^ to TP enabled optical control of microtubule stability. However, these small molecule-based approaches do not generate and control the microtubule superstructures. In contrast, the generation of microtubule superstructures, such as bundles and doublets, was observed by binding of TP-fused tetrameric fluorescence protein, Azami-Green (TP-AG), to the outer surface of microtubules.^23^ Bundle structures were formed by crosslinking of microtubules by tetrameric TP-AG, and doublet microtubules were formed by recruiting free tubulin to the TP moiety of TP-AG-bound microtubules. This study inspired us to hypothesize that a photo-switchable tetrameric protein that converts reversibly between monomeric and tetrameric states by light irradiation can photocontrol microtubule superstructures.

In this report, we focused on a photo-switchable protein, Dronpa, which was developed by Miyawaki et al.,^24,25^ for optical control of microtubule accumulation and dispersion. Tetrameric Dronpa fluoresces green and converts to the monomeric state upon irradiation at 500 nm. In contrast, monomeric Dronpa has no fluorescence and converts to the tetrameric state by 400 nm light irradiation (Figure 1a). In this study, we used the photo-switching of Dronpa for optical control of microtubule assemblies by constructing TP-fused Dronpa construct on its C-terminus (TP-Dronpa). Tetrameric TP-Dronpa facilitated the accumulation of microtubules by crosslinking microtubules with each other, whereas monomeric TP-Dronpa dispersed the microtubule superstructures. Optical control of microtubule superstructures was achieved by switching between the monomeric and tetrameric states of TP-Dronpa.

**Figure 1.**
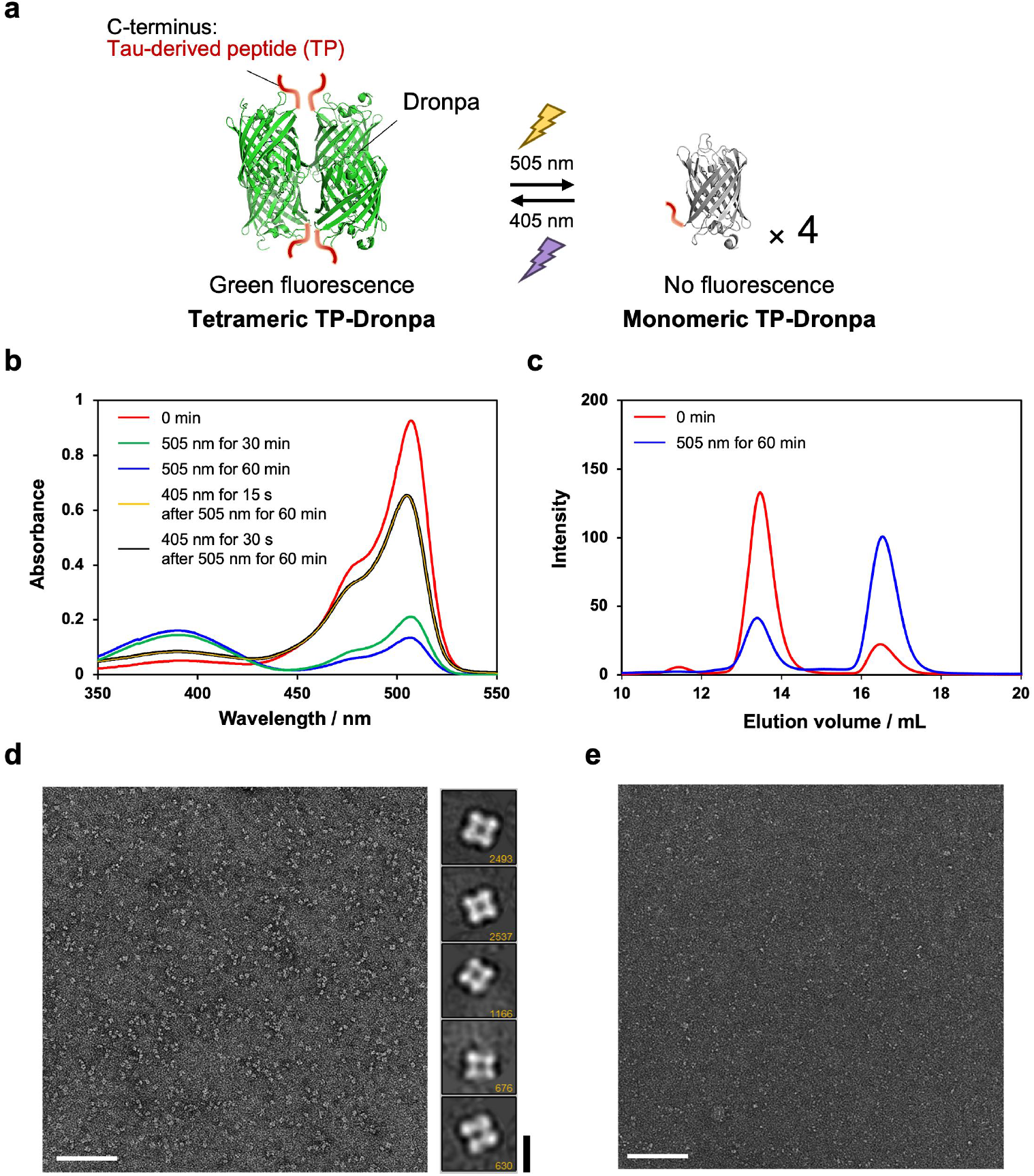
(a) Photo-conversion of the association state of Tau-derived peptide (TP)-fused Dronpa (TP-Dronpa). (b) Light irradiation-dependent changes in TP-Dronpa absorbance. The UV-vis spectra of the initial state (red), with 505 nm light irradiation for 30 min (green) or 60 min (blue), 405 nm light irradiation for 15 s (yellow) or 30 s (black) after 505 nm light irradiation for 60 min at 25 °C. Preparation concentration: 11.3 μM TP-Dronpa. Concentrations of TP-Dronpa are defined as the monomer concentrations. (c) Size exclusion chromatography (SEC) analysis of TP-Dronpa. The initial state of TP-Dronpa (red) and after 505 nm irradiation for 60 min (blue). (d and e) Negative staining transmission electron microscopy (TEM) images of a tetrameric fraction of TP-Dronpa (d, left) and a monomeric fraction of TP-Dronpa (e). Class averages of TP-Dronpa showing clear tetrameric structures by single particle analysis ((d), right). The number of particles is indicated in each image. Scale bars, 100 nm for (d) (left) and (e), 10 nm for (d) (right).

## RESULTS

### Design and Binding Analysis of TP-Dronpa to Microtubules

We designed the TP-Dronpa construct containing the Dronpa145N sequence,^26^ a Dronpa mutant with the K145N mutation, fused with an N-terminal His-tag and C-terminal TP (Figure S1). This construct was used because the previously designed TP-AG with the same positions of the His-tag and TP generated microtubule superstructures.^23^ The plasmid construct was transformed into *E. coli* BL21 and expressed TP-Dronpa was purified by Ni-NTA affinity chromatography (Figure S2). Dronpa without the TP moiety and TP-fused pdDronpa1.2 (TP-pdDronpa) that switches dimeric and monomeric states by light irradiation^27^ were prepared using the same construction approach (Figure S2). UV-vis spectra of TP-Dronpa were recorded to evaluate the conversion of the tetrameric/monomeric states by light irradiation (Figure 1a). The initial state of TP-Dronpa was tetrameric, as defined by the prominent peak at 507 nm and a weak peak at 388 nm, as reported previously (Figure 1b).^24^ Most of the tetrameric state was converted to the monomeric state by irradiating with 505 nm light (130 mW/cm^2^) for 1 h, as observed by an increase in absorption around 400 nm and a decrease in absorption around 500 nm. Conversely, most monomeric TP-Dronpa converted to the tetrameric state by 405 nm light (113 mW/cm^2^) irradiation for 30 s (Figure 1b). Thus, the structural change of TP-Dronpa was reversible. These results align with the reported photoresponsivity of Dronpa.^24^ Size exclusion chromatography (SEC) results also showed that TP-Dronpa is converted from tetramer to monomer by 505 nm light irradiation, with an elution shift from 13.5 mL to 16.5 mL (Figure 1c). The initial state of TP-pdDronpa, which forms a dimer, gave rise to an SEC peak with an elution volume of 14.9 mL (Figure S3), indicating that the peaks of TP-Dronpa at 13.5 and 16.5 mL are tetramer and monomer states, respectively. The existence ratio of TP-Dronpa tetramers and monomers was calculated using the SEC peak areas. For the initial state, 85% of TP-Dronpa formed the tetramer, and 15% of TP-Dronpa formed the monomer. After light irradiation at 505 nm for 1 h, 30% of TP-Dronpa was tetrameric, and 70% of TP-Dronpa was monomeric. The 13.5 mL and 16.5 mL peak fractions were negatively stained and observed by transmission electron microscopy (TEM) (Figure 1d,e). The 13.5 mL peak fraction sample showed unambiguous tetrameric structures with diameters of ∼10 nm, which was also confirmed by single particle analysis. In contrast, the 16.5 mL peak fraction sample showed smaller particles with ∼5 nm diameters corresponding to the TP-Dronpa monomer.

Next, the binding of tetrameric TP-Dronpa to microtubules was evaluated by confocal laser scanning microscopy (CLSM). Tetramethylrhodamine (TMR)-labeled tubulin was polymerized by guanosine-5’-[(α,β)-methyleno]triphosphate (GMPCPP), a GTP analog used to form stable microtubules. TP-Dronpa was incubated with the TMR-labeled microtubules, and CLSM was performed. Co-localization of TMR and Dronpa fluorescence on microtubules was observed (Figure 2a, top left), indicating that TP-Dronpa bound to microtubules. In contrast, tetrameric Dronpa without the TP moiety showed weaker binding to microtubules than TP-Dronpa (Figure S4), indicating that the TP sequence is essential for microtubule binding by TP-Dronpa. Additionally, the binding of dimeric TP-pdDronpa to microtubules was weak (Figure S5). These results indicate that the differences between TP-Dronpa and TP-pdDronpa, such as the number of TPs, size and surface charge (TP-Dronpa has a more negative charge than TP-pdDronpa), affect microtubule binding. The binding site of TP-Dronpa to microtubules was then estimated using subtilisin, an enzyme that selectively digests the C-terminal tails of tubulin located on the outer surface of microtubules.^23,28^ When the microtubules were prepared using subtilisin-treated tubulin, the fluorescence of tetrameric TP-Dronpa localized on the microtubules decreased dramatically (Figure 2a, bottom left). This result indicates that TP-Dronpa mainly binds to the C-terminus of tubulin, the outer surface of microtubules. The binding property of monomeric and tetrameric TP-Dronpa to microtubules was evaluated by a co-sedimentation assay (Figure S6). Free TP-Dronpa and TP-Dronpa bound to microtubules were separated by ultracentrifugation and quantified by SDS-PAGE.^23^ The binding ratio of TP-Dronpa to microtubules estimated in this assay showed that the binding affinity of tetrameric TP-Dronpa to microtubules (*K*_d_ = 9.7 μM) was stronger than that of monomeric TP-Dronpa (*K*_d_ = 51 μM), possibly because of multivalent binding. The binding affinity of tetrameric TP-Dronpa was similar to the previously reported affinity of TP-AG to microtubules (*K*_d_ = 11 μM).^23^ Additionally, the TP-Dronpa-tubulin complex was observed by TEM by incubating TP-Dronpa with tubulin (Figure 2b-d). This result indicates that TP-Dronpa binds to tubulin and microtubules. Because the recruitment of tubulin to TP-AG on microtubules is important for generating microtubule superstructures,^23^ tubulin binding to TP-Dronpa should aid the construction of superstructures.

**Figure 2.**
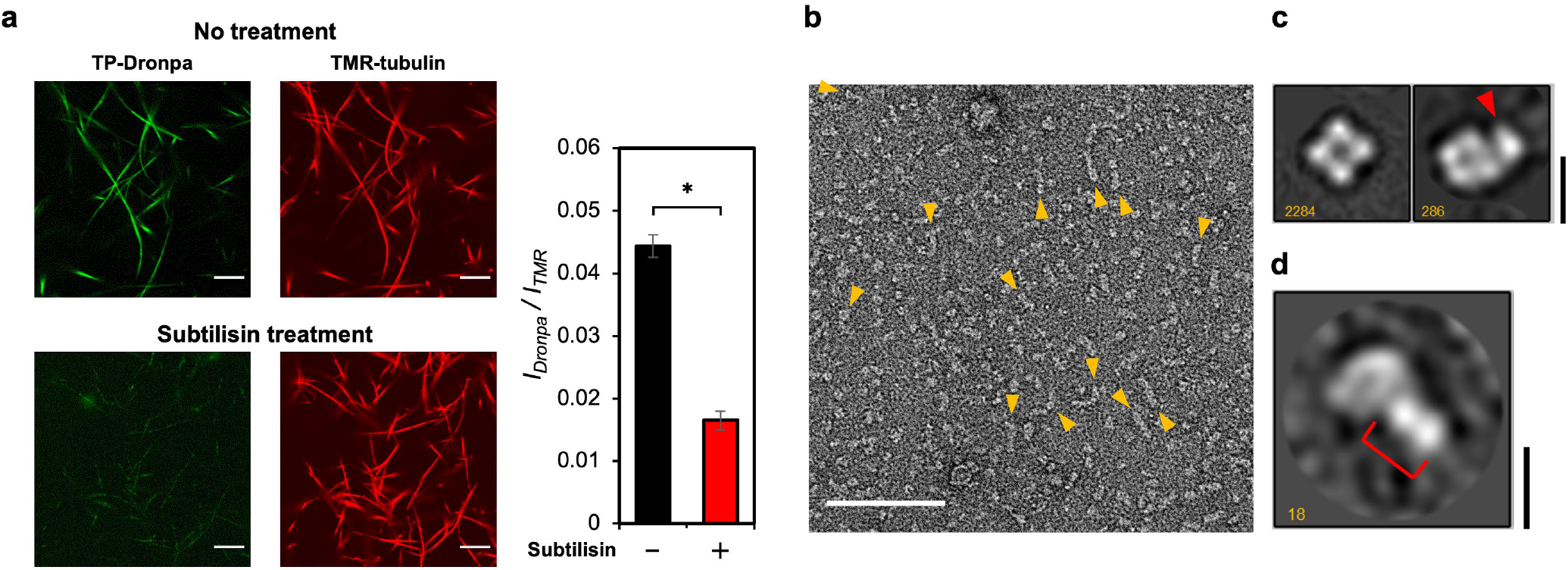
(a) Confocal laser scanning microscopy (CLSM) images of tetrameric TP-Dronpa-incorporated microtubules using intact (top panel) or subtilisin-treated tubulin (bottom panel). Preparation concentrations: 16 μM tubulin, 4 μM TMR-tubulin and 10 μM TP-Dronpa. Scale bars, 10 μm. The *I*_Dronpa_*/I*_TMR_ ratio for each microtubule sample was determined from the CLSM images (right). Error bars represent the standard error of the mean (*N* = 25). \**P* < 0.0001, two-tailed Student’s *t*-test. (b–d) Analysis of the tetrameric TP-Dronpa-tubulin complex by negative staining TEM. (b) A typical micrograph of tetrameric TP-Dronpa incubated with tubulin. Clear interactions are indicated by the orange arrowheads, with tubulin extending from TP-Dronpa to form filamentous polymers. Preparation concentrations: 20 μM TP-Dronpa and 20 μM tubulin. Scale bar, 100 nm. (c, d) Single particle analysis of tetrameric TP-Dronpa in the presence of tubulin. (c) Typical 2D class averages without (left) and with (right) additional density (red arrowhead) after auto-picking. Scale bar, 10 nm. (d) A 2D class average of tetrameric TP-Dronpa binding to tubulin after manual picking. The number of particles is indicated in each image. Scale bar, 10 nm. The red bracket indicates 8-nm additional density.

### Microtubule Accumulation Induced by TP-Dronpa

We next evaluated the effect of TP-Dronpa on the motility of microtubules driven by ATP hydrolysis on a kinesin-coated glass substrate (motility assay). Before adding ATP, the microtubules with tetrameric TP-Dronpa appeared as thick bundle structures, whereas the microtubules with monomeric TP-Dronpa were separated and not bundled (Figure 3a). These results indicate tetrameric TP-Dronpa tethered microtubules by the four TPs exposed on the Dronpa scaffold, whereas crosslinking by monomeric TP-Dronpa to microtubules was not observed. The concentration dependence of TP-Dronpa showed that 0.5 equivalent of TP-Dronpa to tubulin was sufficient to induce the formation of the bundled microtubules (Figure S7). Adding ATP to microtubule bundles with tetrameric TP-Dronpa induced conversion into radially moving aster-like structures and subsequent dissociation (Figure 3b,c and Movie S1). In contrast, microtubules with monomeric TP-Dronpa moved individually like normal microtubules (Movie S2). The conversion of the bundles into aster-like structures may be caused by the random association of microtubule bundles with tetrameric TP-Dronpa in a parallel and anti-parallel manner, followed by dissociation such as asters by the moving of kinesin. These results showed completely different microtubule assembly structures by tetrameric and monomeric TP-Dronpa.

**Figure 3.**
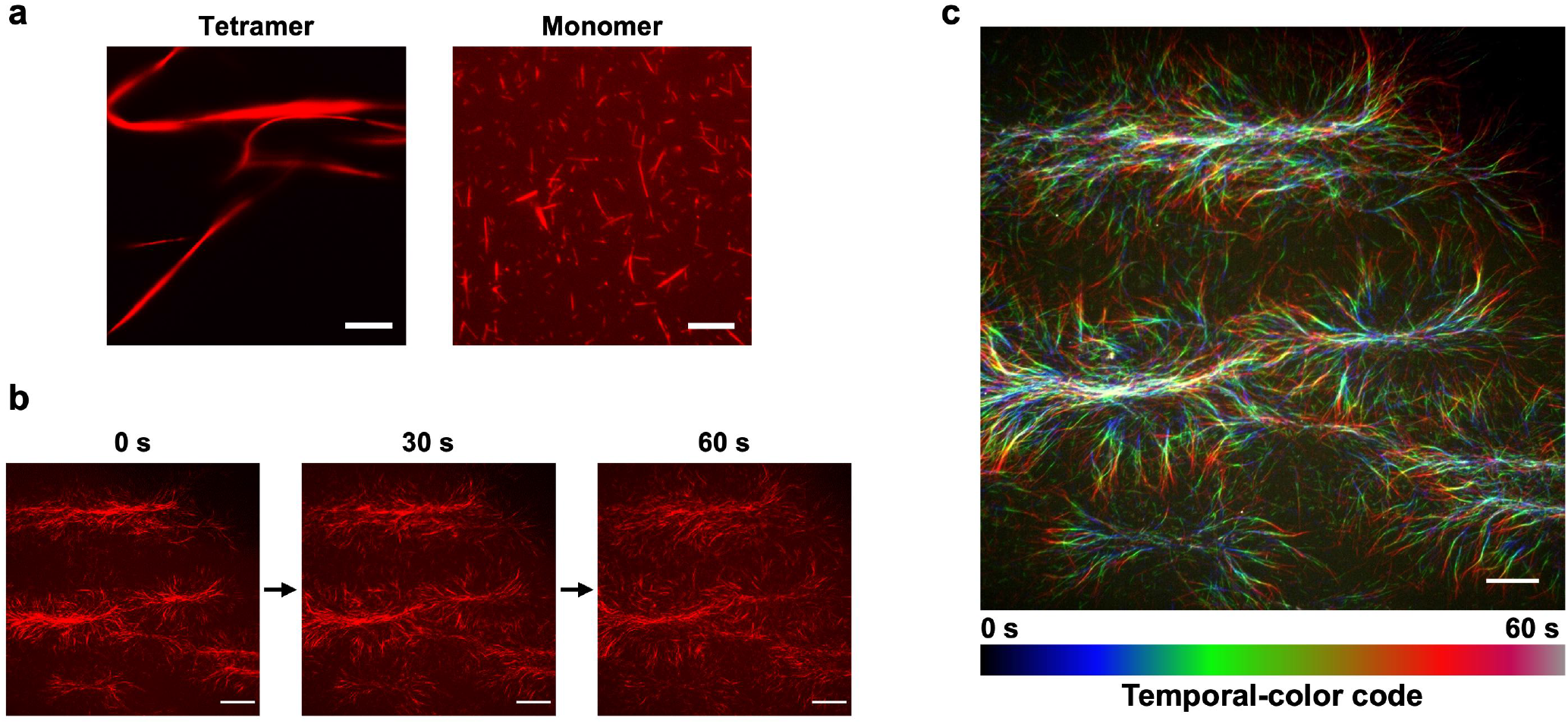
(a) CLSM images of microtubules incorporated with tetrameric TP-Dronpa or monomeric TP-Dronpa. Preparation concentrations: 16 μM tubulin, 4 μM TMR-tubulin and 10 μM TP-Dronpa. Scale bars, 10 μm. (b) Time-lapse images showing the dissociation of bundled microtubules incorporated with tetrameric TP-Dronpa into aster-like structures on the kinesin-coated substrate. Scale bars, 30 μm. (c) A fluorescence microscopy image of moving aster structures in Figure 3b using a temporal-color code. The time series images were color-coded and superimposed to visualize the movement of microtubules. The color scales are shown at the bottom of the image. Scale bar, 20 μm.

### TEM Observation of Microtubule Superstructures

TP-Dronpa-incorporated microtubule structures were observed by TEM. Initially, bundled microtubules with around 200 nm widths were observed by treatment with tetrameric TP-Dronpa (Figure 4a, left). In addition to bundled structures and normal singlet microtubules with 25 nm diameter, doublet-like microtubules consisting of two microtubules with 25 nm (A-tubules) and 15 nm (B-tubules) diameters, typically observed in flagella and cilia,^8^ were observed (Figures 4a, right and S8). The formation of doublet microtubules has been observed previously by treatment of tetrameric TP-AG.^23^ Similar to TP-AG, tetrameric TP-Dronpa bound to the outer surface of microtubules and recruited free tubulin by binding the TP moiety of TP-Dronpa (Figure 2d) to induce the formation of doublet microtubules. In contrast, bundled and doublet microtubules were not observed for monomeric TP-Dronpa (Figure 4b). These results indicate that tetrameric TP-Dronpa induced the formation of microtubule bundles and doublet structures, whereas monomeric TP-Dronpa showed no such effects, supporting the fluorescent microscopy results (Figure 3a).

**Figure 4.**
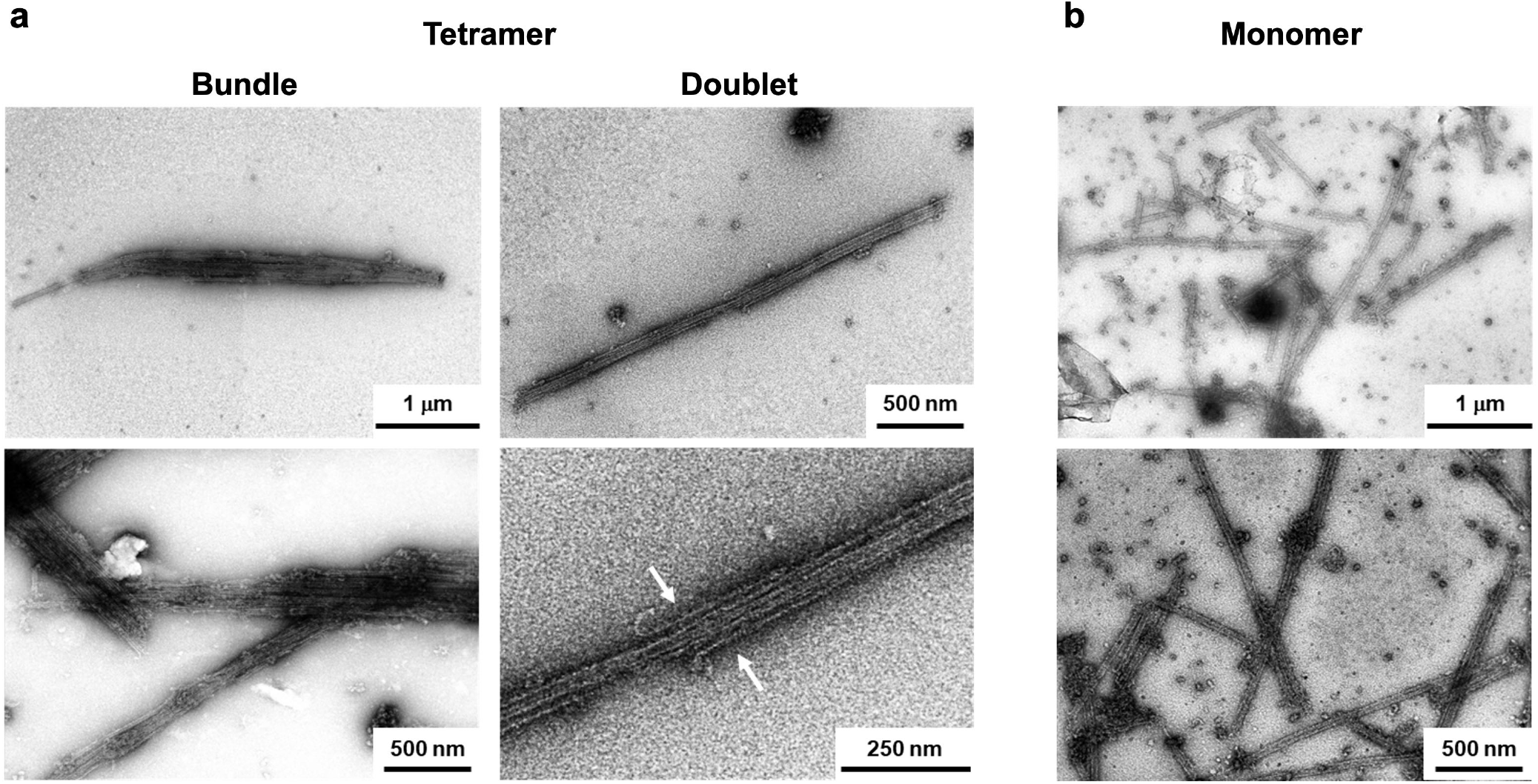
TEM images of TP-Dronpa-incorporated microtubules. Preparation concentrations: 16 μM tubulin, 4 μM TMR-tubulin and 10 μM TP-Dronpa. (a) Bundled microtubules (left) and doublet microtubules (right) bound with tetrameric TP-Dronpa. The B-tubules of doublet microtubules were observed (white arrow). (c) Dispersed microtubules bound with monomeric TP-Dronpa.

### Optical Control of Microtubule Accumulation and Dispersion

Optical control of microtubule accumulation and dispersion was demonstrated by converting monomeric and tetrameric states of TP-Dronpa. Tetrameric TP-Dronpa-incorporated microtubules were prepared, and irradiation with 505 nm light for 1 h was performed to convert tetrameric TP-Dronpa to the monomeric state. Light irradiation converted the bundled microtubules into dissociated and dispersed microtubules (Figure 5a). The fluorescence of tetrameric TP-Dronpa on the microtubules was also observed to decrease following 505 nm light irradiation, indicating the conversion of tetrameric TP-Dronpa to the monomer state. The dispersed microtubules moved individually on the kinesin substrate, as seen for normal microtubules (Movie S3). When monomeric TP-Dronpa-incorporated microtubules were irradiated with 405 nm light for 30 s to convert TP-Dronpa from the monomeric to tetrameric state, the dispersed microtubules accumulated into bundles, with partial recovery of the fluorescence of tetrameric TP-Dronpa (Figure 5b). These bundled microtubules converted aster-like structures and dissociated by the moving of kinesin (Movie S4), as observed in the microtubules with tetrameric TP-Dronpa (Figure 3). These results indicate that microtubule accumulating structures were modulated by the photo-converting tetrameric and monomeric states of TP-Dronpa. The photo-conversion ratio of monomeric TP-Dronpa bound to microtubules into the tetrameric state was estimated by the fluorescence intensity from tetrameric TP-Dronpa by comparing before and after 405 nm light irradiation and after removal of free TP-Dronpa from solutions by ultracentrifugation (Figure S9). The result showed that ∼18% of monomeric TP-Dronpa bound to microtubules was converted to the tetrameric state by 405 nm light irradiation. Thus, the photocontrol of microtubule accumulating structures was achieved by reversible conversion of the monomeric and tetrameric states of TP-Dronpa on microtubules.

**Figure 5.**
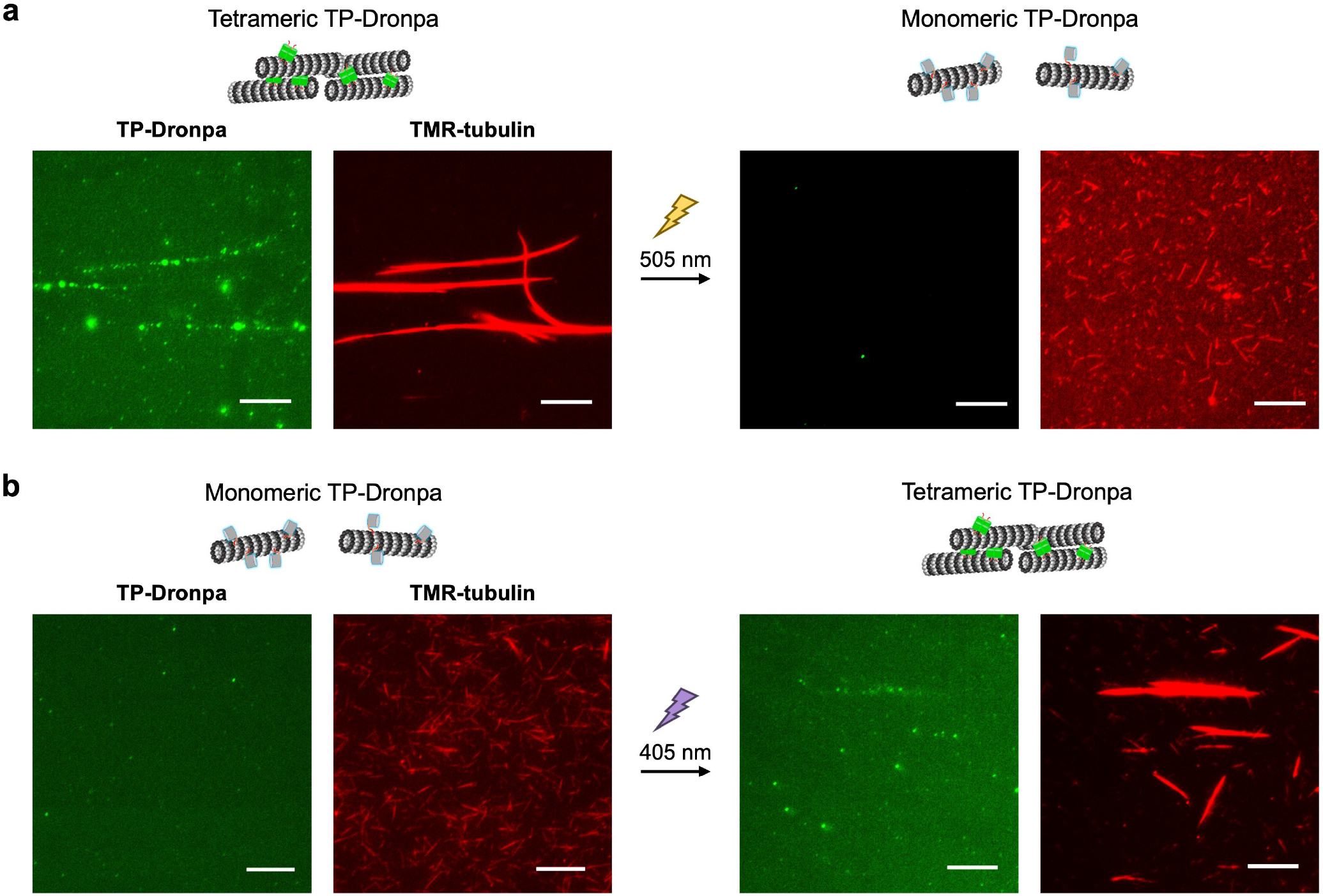
Optical control of microtubule accumulation and dispersion by TP-Dronpa. (a) Dissociation of microtubule assembly by 505 nm light irradiation for 1 h to microtubules incorporated with tetrameric TP-Dronpa. (b) Accumulation of microtubules by 405 nm light irradiation for 30 s to microtubules incorporated with monomeric TP-Dronpa. Preparation concentrations: 16 μM tubulin, 4 μM TMR-tubulin and 10 μM TP-Dronpa. Scale bars, 10 μm.

### Optical Control of Microtubule Accumulation and Swarming

To achieve temporal accumulation and “swarming” movement of microtubules by TP-Dronpa, we used methylcellulose, which is frequently used as a depletion agent to promote the association of microtubules in the motility assay.^29^ Under depleting conditions using methylcellulose, microtubule accumulation and swarming movement in a parallel direction are promoted. In our system, methylcellulose was used to increase the affinity of TP-Dronpa to microtubules during the motility assay for optical control of microtubule accumulation and swarming. When tetrameric TP-Dronpa, ATP and 0.2% methylcellulose were added to microtubules fixed on the kinesin substrate, accumulation of microtubules and swarming movement were observed (Figure 6a, Movie S5). Under these conditions, tetrameric TP-Dronpa possibly crosslinked microtubules, which moved in the same direction to form the accumulated structures in parallel. In contrast, microtubules moved individually using monomeric TP-Dronpa (Figure 6a, Movie S5). The velocity of these microtubules was 0.20 ± 0.009 μm/s for the monomeric TP-Dronpa-incorporated microtubules and 0.077 ± 0.020 μm/s for the tetrameric TP-Dronpa-incorporated microtubules (Figure S10). The reduced velocity of microtubules with tetrameric TP-Dronpa may arise from the suppression of microtubule movement because of induced anti-parallel crosslinking of moving microtubules by a fraction of tetrameric TP-Dronpa.

**Figure 6.**
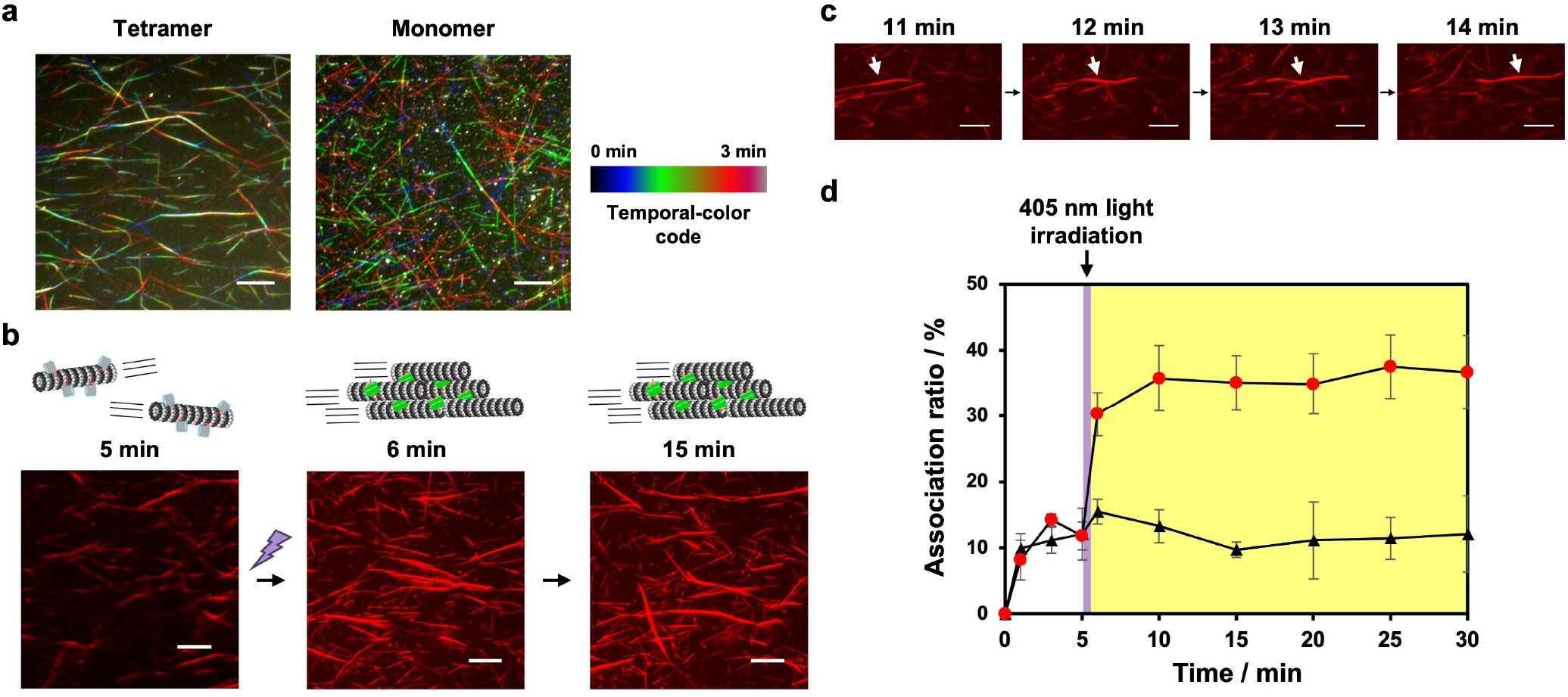
Photocontrol of motile TP-Dronpa-incorporated microtubule assembly under depleting conditions. (a) Fluorescence microscopy images of microtubules incorporated with tetrameric TP-Dronpa or monomeric TP-Dronpa on a kinesin substrate in the presence of 0.2% methylcellulose. Preparation concentrations: 16 μM tubulin, 4 μM TMR-tubulin and 10 μM TP-Dronpa. Scale bars, 10 μm. (b) Time-lapse images showing the accumulation of monomeric TP-Dronpa-incorporated microtubules by 405 nm light irradiation for 30 s at 5 min. Preparation concentrations: 1 μM tubulin, 1 μM TMR-tubulin and 5 μM TP-Dronpa. Scale bars, 10 μm. (c) Representative time-lapse fluorescence microscopy images showing the formation of a bundled microtubule. Scale bars, 10 μm. (d) Change of the association ratio of microtubules over time with 405 nm light irradiation at 5 min (red dot) and without light irradiation (black triangle). Error bars represent the standard deviation of the mean (*N* = 3).

Accumulation and swarming of microtubules were observed by treating tetrameric TP-Dronpa, whereas monomeric TP-Dronpa showed no such effect (Figure 6a), and therefore temporal optical control of microtubule accumulation was attempted. The movement of microtubules with monomeric TP-Dronpa was observed for 5 min, and then the sample was irradiated at 405 nm for 30 s to convert the monomeric form to the tetrameric state. Irradiation at 405 nm induced the accumulation of microtubules and swarming movement (Figure 6b,c, Movie S6). The association ratio of microtubules increased drastically after 405 nm light irradiation (Figure 6d). The photoconverted tetrameric TP-Dronpa on microtubules probably crosslinked microtubules in a parallel manner. These results show the optical control of microtubule accumulation and swarming by conversion of monomeric and tetrameric states of TP-Dronpa bound with microtubules.

## DISCUSSION

In this report, control of accumulation and dispersion of microtubules was achieved by photoconverting the monomeric/tetrameric states of TP-Dronpa. Modifying photoresponsive DNAs to microtubules was used previously to control the accumulation/dispersion of microtubules by light irradiation.^15–17^ However, this system required direct conjugation of DNA to microtubules and could not be used to control intact microtubules. Moreover, the accumulated structures were restricted to swarming microtubules. In this study, various microtubule superstructures were generated and photocontrolled by TP-Dronpa without directly modifying microtubules. Interestingly, random accumulation (parallel and anti-parallel) of microtubules and conversion to aster-like structures were observed by tetrameric TP-Dronpa, whereas swarming microtubules in a parallel direction were generated on a kinesin substrate with depletion force. In addition, our system uses a fluorescence switchable protein that will facilitate tracking microtubule accumulation and dispersion by fluorescent imaging. Moreover, this protein-based accumulation/dispersion system can be used to model natural systems forming microtubule superstructures.

## CONCLUSION

Optical control of microtubule accumulation and dispersion was achieved by Tau-derived peptide-fused photo-switchable Dronpa. TP-Dronpa bound to the outer surface of microtubules and formed bundling, doublet and aster-like microtubule superstructures in the tetrameric state. Optical control of these microtubule superstructures was achieved by photoconversion between the monomeric and tetrameric states of TP-Dronpa. This work demonstrates protein-based optical control of intact microtubules mimicking the formation of microtubule superstructures in nature. In the future, the TP-Dronpa system, a genetically encodable protein system, will be applied to manipulate cell shape and migration direction by optical control of intracellular microtubule accumulation and will be used in nanomaterial research, such as the development of active matter and molecular robotics.

## ACKNOWLEDGMENTS

We thank a technical staff Ms. Noriko Matsuura from Tottori University for the help of protein purification and Bio-Electron Microscopy Facility of ShanghaiTech University for TEM imaging. This study was financially supported by FOREST Program (JPMJFR2034) and ACT-X (JPMJAX2012) from the Japan Science and Technology Agency (JST) and JSPS KAKENHI Grant number JP23K04931 and JP24H01721 in a Grant-in-Aid for Transformative Research Areas “Materials Science of Meso-Hierarchy” from the Japan Society for the Promotion of Science (JSPS) to H.I., and by JST, PRESTO (JPMJPR20E1) to M.I. We thank Edanz (https://jp.edanz.com/ac) for editing a draft of this manuscript.

