## Supporting Information for "Optical Control of Microtubule Accumulation and Dispersion by Tau-Derived Peptide-Fused Photo-Responsive Protein"

### Experimental Section

#### Equipment and materials.

UV-vis spectra were obtained using a Jasco V-630. Fluorescence spectra were obtained using a Jasco FP-8200. Ultracentrifugation was performed using an Optima MAX-TL ultracentrifuge (Beckman Coulter) using TLA 120.2 rotor. Tubulin was purified from porcine brain by a reported procedure.[1] Recombinant kinesin-1 consisting of the first 573 amino acid residues of human kinesin-1 was prepared according to a reported procedure.[2] SEC was performed using an ÄKTA™ avant (Cytiva). In the SDS-PAGE analysis, the proteins were mixed with 2× Laemmli buffer, heated at 95 °C for 5 min, and then loaded onto SDS-PAGE gel (15% acrylamide) and electrophoresis was carried out at 200 V (constant voltage) in a buffer. TEM images were obtained using JEOL JEM 1400 Plus and Talos L120C TEM (Thermo Fisher Scientific). The reagents were purchased from Nacalai Tesque, Inc., COSMO BIO Co. Ltd., Tokyo Chemical Industry Co., Dojindo Laboratories Co. Ltd., Wako Pure Chemical Industries, and Sigma-Aldrich and used without further purification. Confocal laser scanning microscopy (CLSM) measurement was carried out using a FluoView FV10i (Olympus). In the motility assay, samples were illuminated with a light-emitting diode light source and visualized using an epi-fluorescence microscope (Eclipse Ti2-E, Nikon) using an oil-coupled Lambda S 60× objective [numerical aperture (NA) 1.4] (Nikon). Concentrations of TP-Dronpa, Dronpa, and TP-pdDronpa are described as monomer concentrations throughout manuscript.

#### Construction of plasmids.

The TP-Dronpa, Dronpa, and TP-pdDronpa expression vectors were constructed on the basis of the pET-29b(+) expression vector (Merck, Darmstadt, Germany). The TP-Dronpa, Dronpa, and TP-pdDronpa genes were synthesized to optimize codon usage for *Escherichia coli*. The open circular pET-29b(+) vector was synthesized by inverse polymerase chain reaction (PCR). PCR was performed in the TaKaRa PCR Thermal Cycler Dice Touch (TaKaRa) using the Tks Gflex DNA Polymerase (TaKaRa). The synthetic genes coding TP-Dronpa, Dronpa, and TP-pdDronpa were ligated into the open circular pET-29b(+) using the In-Fusion HD Cloning Kit (TaKaRa) to generate the recombinant protein expression vectors. The plasmids were sequenced by Eurofins Genomics.

#### Expression and purification of proteins.

The pET-29b(+) vectors coding TP-Dronpa, Dronpa, and TP-pdDronpa were transformed into *E. coli* ClearColi BL21(DE3) strain (Biosearch Technologies) by a heat-shock procedure. Bacterial cells were spread on Luria-Bertani– Ampicillin (100 μg/mL) (LBA) agar and grown overnight at

37 °C. A single transformant colony was grown in LBA medium at 37 °C overnight. The culture was diluted 100-fold by addition to fresh LBA medium and grown at 37 °C until an absorbance of 0.5 was noted at 600 nm (mid-logarithmic phase), and then the culture was incubated with 0.1 mM isopropyl β-D-1-thiogalactopyranoside at 20 °C. After 18 h of incubation, cells were harvested by centrifugation at 8,000 rpm for 10 min. The cell pellets were suspended in Ni-affinity binding buffer [50 mM tris-HCl (pH 7.4), 150 mM NaCl, and 20 mM imidazole] on ice. The cells were lysed by sonication. After centrifugation at 13,000 rpm for 10 min, the supernatant was loaded onto 1 mL of Ni-affinity column (Cytiva). After washing with the same buffer and the storage buffer [50 mM tris-HCl (pH 8.0) and 300 mM NaCl], the protein was eluted from the column using the Ni-affinity elution buffer [50 mM tris-HCl (pH 8.0), 300 mM NaCl, and 250 mM imidazole]. The eluted sample was dialyzed against the storage buffer [Spectra/por7; cutoff molecular weight (Mw), 8 kDa; Spectrum Laboratories Inc.] at 4 °C. The purity of proteins was evaluated by SDS-PAGE (Figure S2). The concentration of proteins was determined using reported molar extinction coefficients.[3]

**UV-vis spectrum measurement.**

LED light (CL-1503 or CL-1501, ASAHI SPECTRA, 405 nm: 113 mW/cm$^2$ or 505 nm: 130 mW/cm$^2$) was irradiated to the solution of 11.3 μM TP-Dronpa in the storage buffer with 505 nm light for 30 min or 1 h at a distance of 9 cm from bottom of Eppendorf tube, and then UV-vis spectra were measured at 25 °C. After that, 405 nm light was irradiated to the solution of TP-Dronpa and then UV-vis spectra were measured at 25 °C.

**SEC analysis.**

For TP-Dronpa, protein sample purified by Ni-NTA purification (8.3 mg/mL) was diluted fivefold with the storage buffer, achieving a final volume of 500 μL. Then the samples with or without radiation with a 505 nm light (CL-1501, ASAHI SPECTRA) were ultracentrifuged (25,000 rpm for 20 min, himac S55A2 rotor), and then applied to a Superdex 200 Increase 10/300 GL column (Cytiva) equilibrated with the storage buffer. For TP-pdDronpa protein, 9.5 mg/mL protein sample after Ni-NTA purification was ultracentrifuged (25,000 rpm for 20 min, himac S55A2 rotor), and then applied to a Superdex 200 Increase 10/300 GL column (Cytiva) equilibrated with the storage buffer. The fractions with target proteins were collected and pooled.

**Preparation of tetramethylrhodamine (TMR)-labeled tubulin (TMR-tubulin).**

TMR-tubulin was prepared using 5-carboxytetramethylrhodamine succinimidyl ester according to the standard procedure.[4] The labeling ratio of TMR to tubulin was determined by measuring the absorbance of the protein and TMR at 280 and 555 nm, respectively.

**CLSM measurements.**

The glass bottom dishes (Matsunami, Osaka, Japan) were coated by 1 mg/mL poly-L-lysine (Mw: 30000–70000, Sigma) at room temperature for 15 min, then removed and dried. The microtubule samples were put on the plate and kept at room temperature for 30 min, then observed by CLSM. TMR-tubulin was excited with 550 nm and observed through a 574 nm emission band-pass filter (Red). Dronpa and pdDronpa were excited with 499 nm and observed through a 520 nm emission band-pass filter (Green).

**Binding analysis of TP-Dronpa to microtubules.**

C-terminal tails of tubulin were removed by the treatment of subtilisin according to the reported procedure with modification.[5] Mixture of tubulin (192 μM) and TMR-tubulin (48 μM) in BRB80 [80 mM PIPES (pH 6.9), 1.0 mM $MgCl_2$, and 1.0 mM EGTA] were treated with subtilisin in a subtilisin:tubulin ratio of 1:500 (w/w) at 4 °C for 30 min. Then the subtilisin activity was inhibited by incubation with 5 mM phenylmethylsulfonyl fluoride at 4 °C for 10 min. After centrifugation at 21,000 rpm at 4 °C for 20 min, the supernatants were collected and used as subtilisin-treated tubulin. Then 2 μL of GMPCPP premix (1 mM GMPCPP and 20 mM $MgCl_2$ in BRB80) was added to 8 μL of the solution containing subtilisin-treated tubulin in BRB80. The mixture was incubated at 37 °C for 30 min in the dark. Then 60 μM TP-Dronpa (2 μL) was added to the mixture and kept at 25 °C for 30 min in the dark (final concentrations: [tubulin] = 19.2 μM, [TMR-tubulin] = 4.8 μM, [TP-Dronpa] = 12 μM, and [GMPCPP] = 0.2 mM). As a control, tubulin without subtilisin treatment was used instead of subtilisin-treated tubulin. Dronpa without TP and TP-pdDronpa were also used for the binding experiments using tubulin without subtilisin treatment. TP-Dronpa and TMR fluorescence intensity per microtubule were measured from the fluorescence images by subtracting the background intensity using ImageJ software. The background-subtracted TP-Dronpa fluorescence intensity and TMR fluorescence intensity for each microtubule ($N = 25$) was calculated from 5 images to estimate the binding of TP-Dronpa to microtubules.

**Co-sedimentation assay.**

Tetrameric TP-Dronpa was used in the initial state of purified TP-Dronpa. Monomeric TP-Dronpa was prepared by 505 nm light irradiation (130 mW/$cm^2$) to tetrameric TP-Dronpa for 1 h at room temperature. GMPCPP premix (6 μL) was mixed with a solution (18 μL) containing tubulin (0, 8.3, 16.7, 33.3, 66.7 μM) in BRB80. The mixture was incubated at 37 °C for 30 min in the dark. Then, tetrameric or monomeric TP-Dronpa (100 μM, 6 μL) was added to the mixture and kept at 25 °C for 30 min in the dark (final concentrations: [tubulin] = 0, 5, 10, 20, 40 μM, [TP-Dronpa] = 20 μM, and [GMPCPP] = 0.2 mM). After ultracentrifugation of the mixture at

50,000 rpm at 37°C for 5 min, the supernatants and the pellets were collected and analyzed by SDS-PAGE. The ratio of TP-Dronpa bound to microtubules ($\theta$) was calculated as the ratio of the density of TP-Dronpa bands in the supernatants and the pellets. The results were treated with the following Hill equation (Eq. 1) to give the dissociation constant ($K_d$) and the Hill coefficient ($n$).

$$\log\left(\frac{\theta}{1-\theta}\right) = n \log\frac{1}{K_d} + n \log[Tubulin] \qquad (1)$$

**Fluorescence spectrum measurement.**

GMPCPP premix (20 μL) was mixed with a solution (60 μL) containing tubulin (33 μM) in BRB80. The mixture was incubated at 37 °C for 30 min in the dark. Then, monomeric TP-Dronpa (105 μM, 20 μL) was added to the mixture and kept at 25 °C for 30 min in the dark (final concentrations: [tubulin] = 20 μM, [TP-Dronpa] = 21 μM, and [GMPCPP] = 0.2 mM). After ultracentrifugation of the mixture at 50,000 rpm at 37 °C for 5 min, the pellets were collected and resuspended with BRB80. Fluorescence was measured by fluorescence spectrophotometer before and after 405 nm light irradiation for 1 min. The conversion ratio was estimated fluorescence derived from tetrameric TP-Dronpa subtracted fluorescence before light irradiation.

**Motility assay.**

GMPCPP premix (2 μL) was mixed with a solution (5 μL) containing tubulin (32 μM) and TMR-tubulin (8 μM) in BRB80. The mixture was incubated at 37 °C for 30 min in the dark. Then TP-Dronpa (34 μM, 3 μL, monomer or tetramer prepared as above) was added to the mixture and kept at 25 °C for 30 min in the dark (final concentrations: [tubulin] = 16 μM, [TMR-tubulin] = 4 μM, [TP-Dronpa] = 10 μM, and [GMPCPP] = 0.2 mM). The samples were diluted 50-fold by BRB80. Flow cells were prepared by making a narrow channel on a 24-mm by 60-mm coverslip covered with a 18-mm by 18-mm coverslip (Matsunami, Osaka, Japan) using double-sided tape as a spacer. First, casein (0.5 mg/mL) in BRB80 was introduced into the flow cells and incubated for 3 min. Then, the solution was exchanged with wash buffer [casein (0.5 mg/mL), D-glucose (4.5 mg/mL), glucose oxidase (50 U/mL), catalase (50 U/mL), 1.0 mM dithiothreitol, and 1.0 mM $MgCl_2$ in BRB80] containing 700 nM kinesin and incubated for 3 min. After washing with wash buffer, the solution was exchanged with microtubule solution and incubated for 3 min. After washing with wash buffer, the solution was exchanged with wash buffer containing 5.0 mM ATP and 1.0 mM Trolox. Then, the motility of microtubules was imaged every 10 s. All the experiments were performed at room temperature.

**Concentration dependence of TP-Dronpa on the bundled structures of microtubules.**

GMPCPP premix (2 μL) was mixed with a solution (5 μL) containing tubulin (32 μM) and tubulin-TMR (8 μM) in BRB80. The mixture was incubated at 37 °C for 30 min in the dark. Then tetrameric TP-Dronpa (3.4, 34, 268 μM, 3 μL) was added to the mixture and kept at 25 °C for 30 min in the dark (final concentrations: [tubulin] = 16 μM, [tubulin-TMR] = 4 μM, [TP-Dronpa] = 1, 10, 80 μM, and [GMPCPP] = 0.2 mM). The samples were diluted 50-fold by BRB80. Flow cells were prepared as above. First, casein (0.5 mg/mL) in BRB80 was introduced into the flow cells and incubated for 3 min. Then, the solution was exchanged with wash buffer [casein (0.5 mg/mL), D-glucose (4.5 mg/mL), glucose oxidase (50 U/mL), catalase (50 U/mL), 1.0 mM dithiothreitol, and 1.0 mM $MgCl_2$ in BRB80] containing 700 nM kinesin and incubated for 3 min. After washing with wash buffer, the solution was exchanged with microtubule solution and incubated for 3 min. After washing with wash buffer, microtubules were imaged by fluorescence microscopy.

**TEM observation of TP-Dronpa and TP-Dronpa incubated with tubulin.**

Monomeric and tetrameric TP-Dronpa fractions after SEC were diluted with the storage buffer (final concentration: [TP-Dronpa] = 15 μg/mL) and used for negative staining TEM. For TP-Dronpa-tubulin complex, tetrameric TP-Dronpa fraction and tubulin were mixed (Final concentrations: [TP-Dronpa] = 20 μM, [Tubulin] = 20 μM), and incubated at 25 °C for 30 min, and the sample was diluted 100 times. Sample solutions (3.5 μL) were applied to carbon-coated Cu-grids (U1013, EM Japan Co.,LTD.) pre-hydrophilized using PELCO easiGlow ™ (TED PELLA, INC.), and incubated for 3 min and the excess solution was removed. Then, grids were flipped and sequentially touched to two drops of Milli-Q water and two drops of 1.5% uranyl acetate, then put on a drop of 1.5% uranyl acetate for 30 s. The remaining solution on the grids was removed by filter paper and dried at room temperature. After that, samples were imaged in a TEM (Talos L120C TEM, Thermo Fisher Scientific) operating at an accelerating voltage of 120 kV. Images were collected for each sample in a pixel size of 1.548 Å/pixel.

**Single particle analysis.**

Obtained TEM images were imported into Relion 3.0. For tetrameric TP-Dronpa sample, particles were picked automatically using the template-free procedure based on a Laplacian-of-Gaussian (LoG) filter in Relion, yielding a total of 33,114 particles from 60 micrographs, the particles were extracted in a box size of 128 pixel and underwent reference-free 2D classification using a 200-Å mask, and classified into 100 classes. Typical classes showing clear tetrameric structure are shown in Figure 1d. For TP-Dronpa-tubulin complex, the particles were first picked automatically using the LoG filter in Relion, yielding a total of 94,112 particles from 161 micrographs. The particles were extracted with a box size of 128 pixels and classified into 100 classes using reference-free 2D classification with a 200-Å mask. Representative class averages are shown in Figure 2c.

Subsequently, a manual picking was conducted to enhance the clarity of the results. 228 particles of tetrameric TP-Dronpa with additional densities were picked by visual inspection and then extracted with a box size of 192 pixels. These particles were then further classified by a 2D classification using a 270-Å mask, segmented into 5 classes. Typical class average is shown in Figure 2d.

**TEM observation of TP-Dronpa-incorporated microtubules.**

Monomeric or tetrameric TP-Dronpa-incorporated microtubules were prepared as above (Final concentrations: [Tubulin] = 16 μM, [TMR-tubulin] = 4 μM, [TP-Dronpa] = 10 μM, [GMPCPP] = 0.2 mM). Sample solution (5 μL) was applied to carbon-coated Cu-grids (Thin Carbon film coated TEM Grids, Science Service) for 1 min and then removed. Then, 16% EM stainer (Nisshin EM Co., Ltd.) applied to the TEM grids for 3 min, and then removed. After the sample-loaded carbon-coated grids were dried in desiccator. After that, microtubules were imaged by TEM (JEOL JEM 1400 Plus) at an accelerating voltage of 80 kV.

**Optical control of accumulation and dispersion of TP-Dronpa-incorporated microtubules.**

Monomeric or tetrameric TP-Dronpa-incorporated microtubules were prepared as above (Final concentrations: [Tubulin] = 16 μM, [TMR-tubulin] = 4 μM, [TP-Dronpa] = 10 μM, [GMPCPP] = 0.2 mM). For conversion of tetrameric TP-Dronpa to monomeric state, 505 nm light (130 $mW/cm^2$) was irradiated to the tetrameric TP-Dronpa-incorporated microtubules for 1 h at room temperature. For conversion of monomeric TP-Dronpa to tetrameric state, 405 nm light (113 $mW/cm^2$) was irradiated to the monomeric TP-Dronpa-incorporated microtubules for 30 s at room temperature. These samples were diluted 50-fold by BRB80. Flow cells were prepared as above. First, casein (0.5 mg/mL) in BRB80 was introduced into the flow cells and incubated for 3 min. Then, the solution was exchanged with wash buffer [casein (0.5 mg/mL), D-glucose (4.5 mg/mL), glucose oxidase (50 U/mL), catalase (50 U/mL), 1.0 mM dithiothreitol, and 1.0 mM $MgCl_2$ in BRB80] containing 700 nM kinesin and incubated for 3 min. After washing with wash buffer, the solution was exchanged with microtubule solution and incubated for 3 min. After washing with wash buffer, microtubules were imaged by fluorescence microscopy.

**Optical control of motile TP-Dronpa-incorporated microtubule assembly.**

Microtubules were prepared as above (Final concentrations: [Tubulin] = 1 μM, [TMR-tubulin] = 1 μM, [GMPCPP] = 0.2 mM). For conversion of tetrameric TP-Dronpa to monomeric state, 505 nm light (130 $mW/cm^2$) was irradiated for 1 h at room temperature. This sample was diluted 2-fold by BRB80. Flow cells were prepared as above. Before imaging by fluorescence microscopy, solution in the flow cell was exchanged with wash buffer containing 5.0 mM ATP, 1.0 mM Trolox,

0.2% methylcellulose and 5 µM of tetrameric or monomeric TP-Dronpa. In the case of irradiation of 405 nm light for tetramerization of TP-Dronpa during time-lapse imaging, the solution in flow cell was exchanged with wash buffer containing 5.0 mM ATP, 1.0 mM Trolox, 0.2% methylcellulose and 5 µM of monomeric TP-Dronpa before imaging, and the light was irradiated for 30 s on the fluorescence microscope. The association ratio at a given time $t$ was determined by counting the number of single microtubules manually and dividing the number at time $t$ by the number present initially ($t$ = 0). The time-dependent association ratio $R(t)$ of TMR-labeled microtubules was determined as following equation (Eq. 2) with $N_0$ = Initial number of single microtubules, $N(t)$ = Number of single microtubules after time $t$. The mean association ratio was obtained from the average of three regions of interest (73.3 µm × 73.3 µm).

$$R(t) = \frac{N_0 - N(t)}{N_0} \qquad (2)$$

**Dronpa (Dronpa145N-TP)**

MHHHHHHGSMSVIKPDMKIKLRMEGAVNGHPFAIEGVGLGKPFEGKQSMDLKVKEGG
PLPFAYDILTTVFCYGNRVFAKYPENIVDYFKQSFPEGYSWERSMNYEDGGICNATNDIT
LDGDCYIYEIRFDGVNFPANGPVMQKRTVKWEPSTENLYVRDGVLKGDVNMALSLEGG
GHYRCDFKTTYKAKKVVQLPDYHFVDHHIEIKSHDKDYSNVNLHEHAEAHSELPRQAK

**TP-Dronpa (Dronpa145N-TP)**

MHHHHHHGSMSVIKPDMKIKLRMEGAVNGHPFAIEGVGLGKPFEGKQSMDLKVKEGG
PLPFAYDILTTVFCYGNRVFAKYPENIVDYFKQSFPEGYSWERSMNYEDGGICNATNDIT
LDGDCYIYEIRFDGVNFPANGPVMQKRTVKWEPSTENLYVRDGVLKGDVNMALSLEGG
GHYRCDFKTTYKAKKVVQLPDYHFVDHHIEIKSHDKDYSNVNLHEHAEAHSELPRQAK
GGGSGGGKKHVPGGGSVQIVYKPVDL

**TP-pdDronpa (pdDronpa1.2-TP)**

MHHHHHHGSMSVIKPDMKIKLRMEGAVNGHPFAIEGVGLGKPFEGKQSIDLKVKEGGP
LPFAYDILTTAFCYGNRVFAKYPENIVDYFKQSFPEGYSWERSMNYEDGGICNATNDITL
DGDCYIYEIRFRGTNFPANGPVMQKRTVKWEPSTENLYVRDGVLKGDVIMALSLEGGG
HYRCDFKTTYKAKKVVQLPDYHFVDHHIEIKSHDKDYSNVNLHEHAEAHSELPRQAK
GGGSGGGKKHVPGGGSVQIVYKPVDL

**Figure S1.** Amino acid sequences of Dronpa, TP-Dronpa, and TP-pdDronpa. The red indicates Dronpa145N, blue indicates TP, and green indicates pdDronpa1.2.

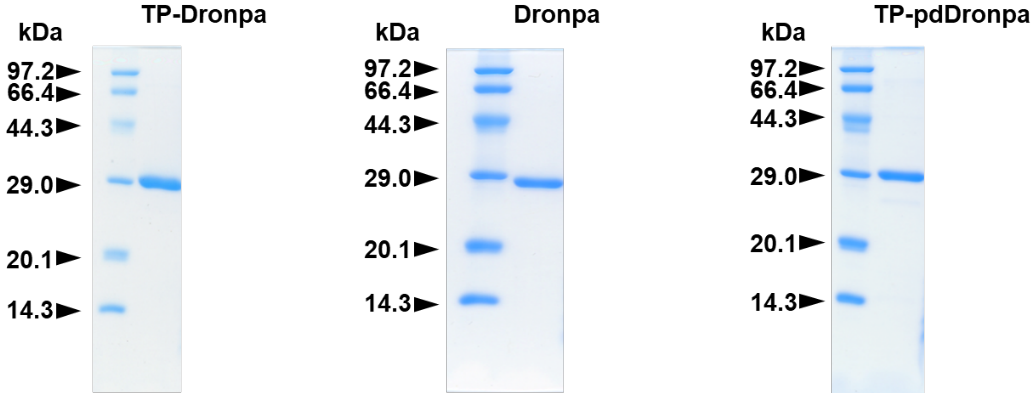


**Figure S2.** SDS-PAGE of TP-Dronpa (29.0 kDa), Dronpa (26.6 kDa) and TP-pdDronpa (29.0 kDa). These results show monomeric states of the proteins.

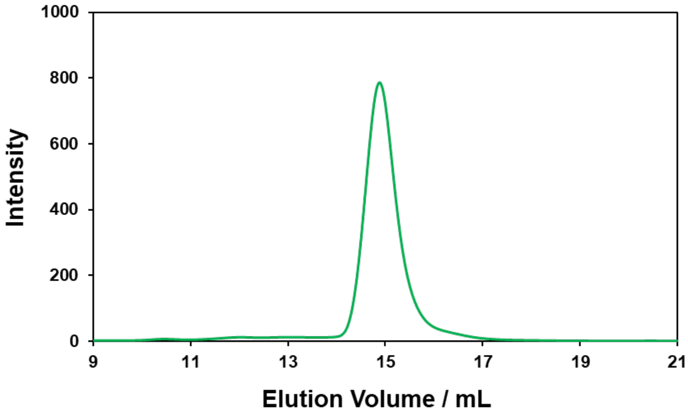


**Figure S3.** SEC analysis of initial state of TP-pdDronpa.

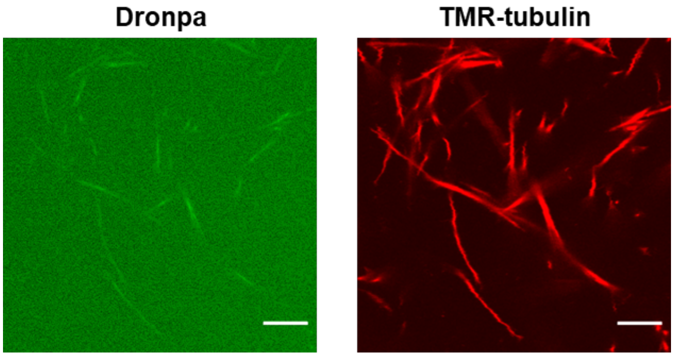


**Figure S4.** CLSM images of microtubules incubated with tetrameric Dronpa. Preparation concentrations: 16 μM tubulin, 4 μM TMR-tubulin and 20 μM Dronpa. Scale bars, 10 μm.

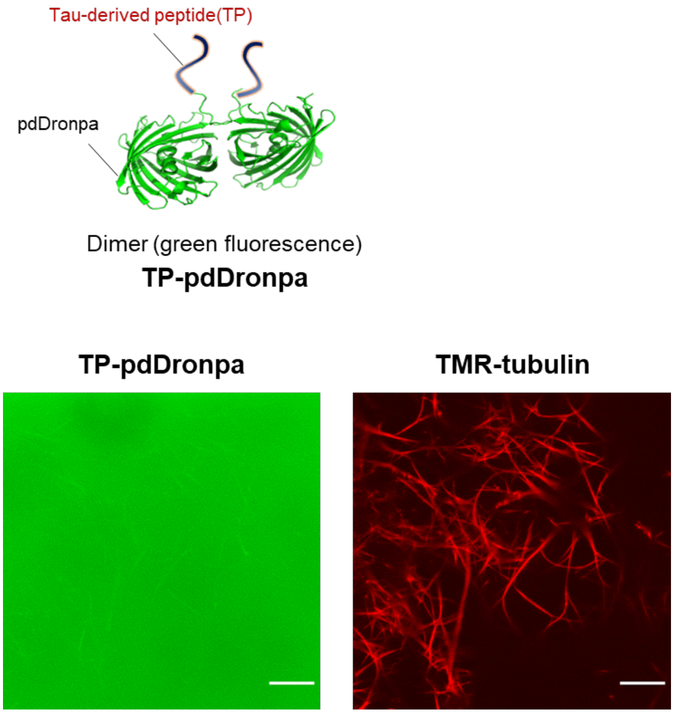


**Figure S5.** CLSM images of microtubules incubated with dimeric TP-pdDronpa. Preparation concentrations: 10 μM tubulin, 10 μM TMR-tubulin and 40 μM TP-pdDronpa. Scale bars, 10 μm.

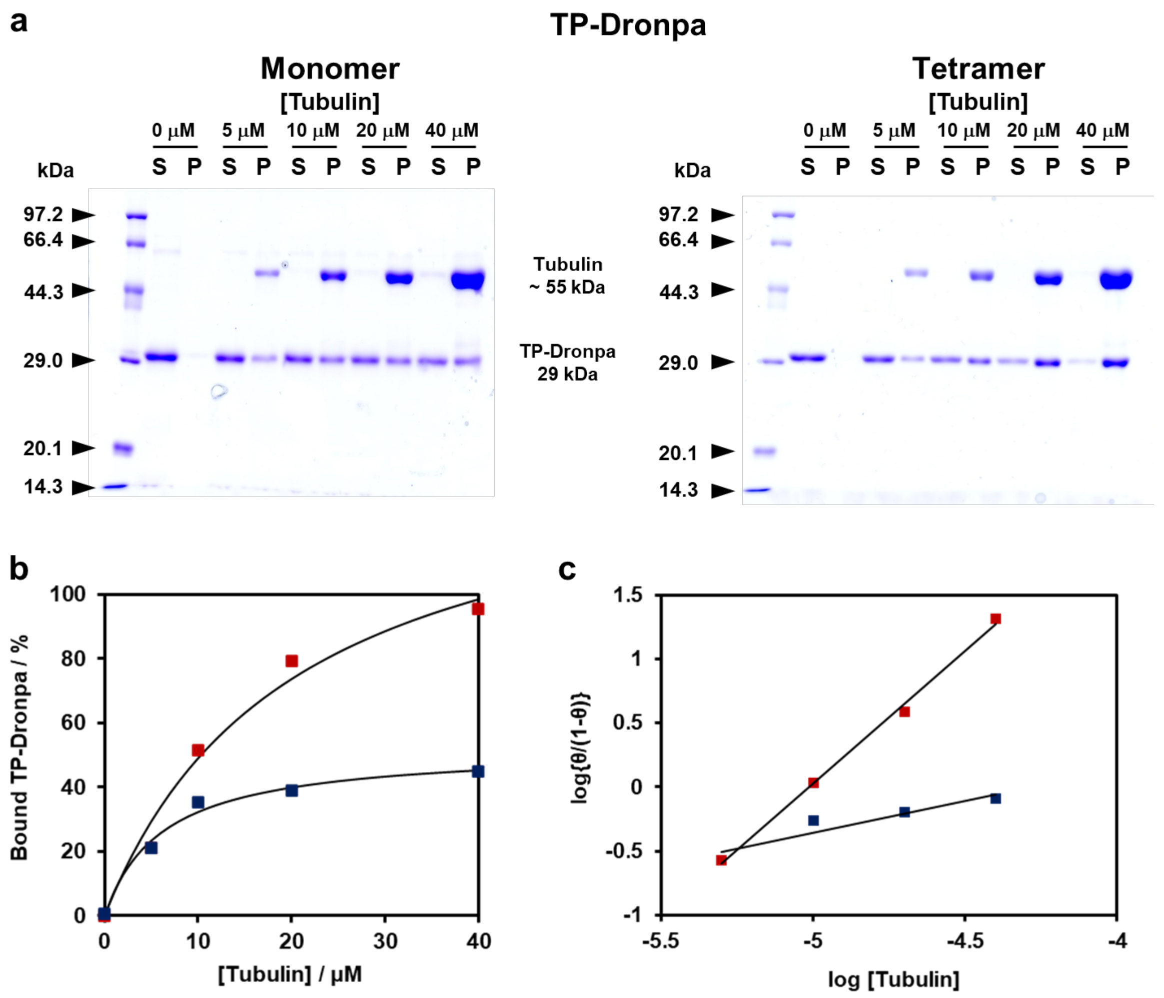


**Figure S6.** Binding of TP-Dronpa to microtubules by co-sedimentation assay. (a) SDS-PAGE of the supernatants (S) and pellets (P) obtained by centrifugation of each sample. (b) Binding ratio of tetrameric or monomeric TP-Dronpa to microtubules and (c) Hill plot. θ indicates the bound ratio of TP-Dronpa to microtubules. Closed square are experimental values (red: tetrameric TP-Dronpa, blue: monomeric TP-Dronpa) and the lines are the fitted values.

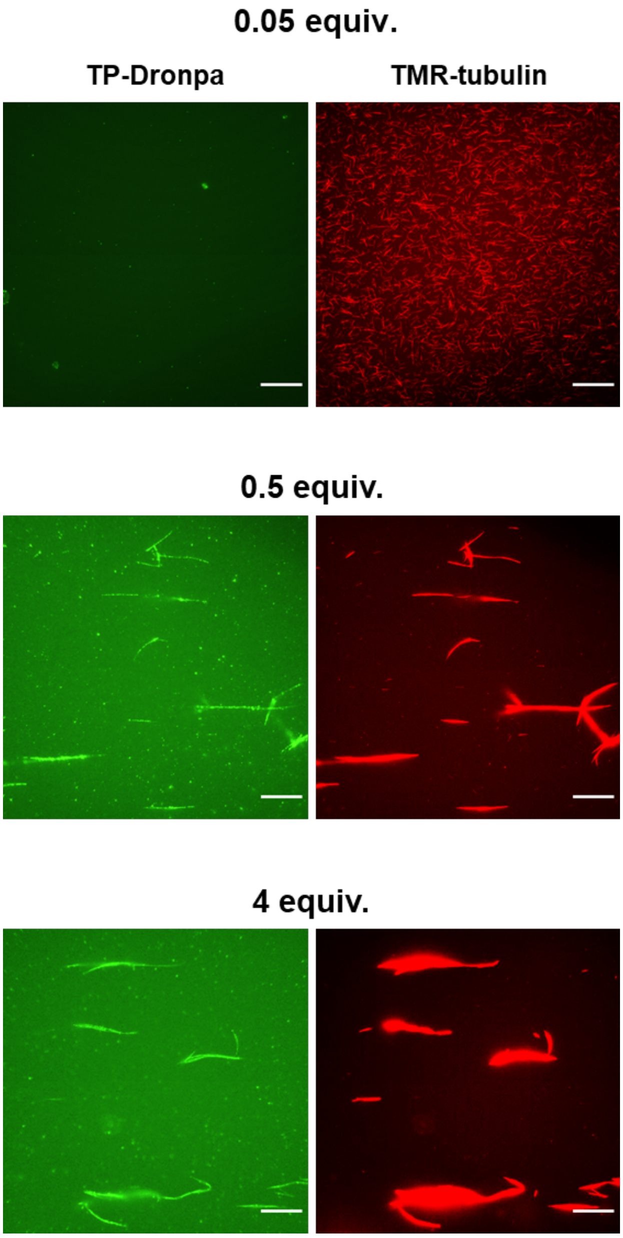


**Figure S7.** Fluorescence microscopy images of microtubules incubated with different equivalents of tetrameric TP-Dronpa to tubulin. Scale bars, 30 μm.

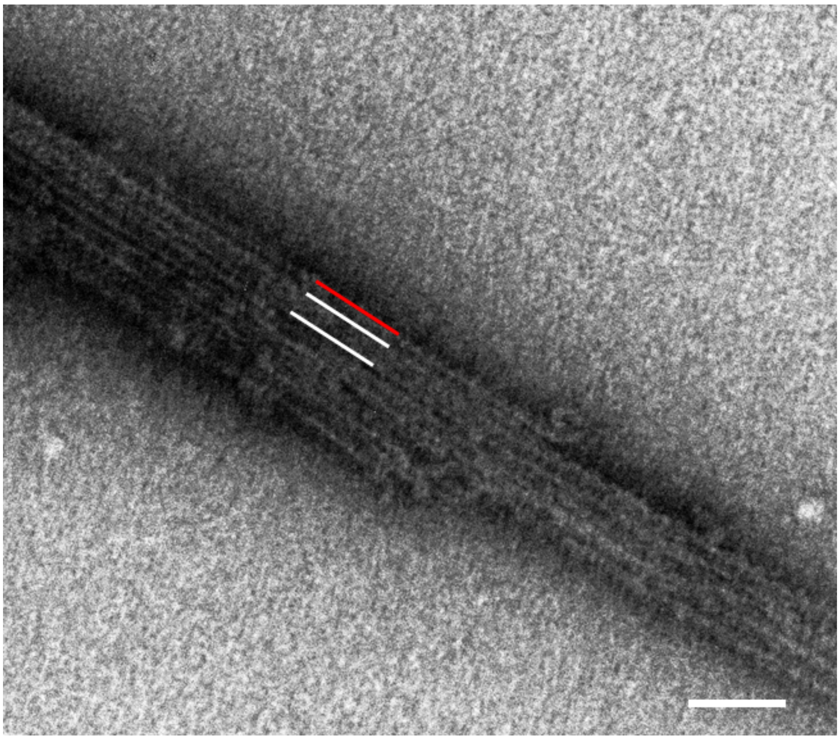

**Figure S8.** Expanded TEM images show doublet-like microtubules in the presence of tetrameric TP-Dronpa. White lines and red line were assumed A-tubule and B-tubule. Scale bar, 100 μm.

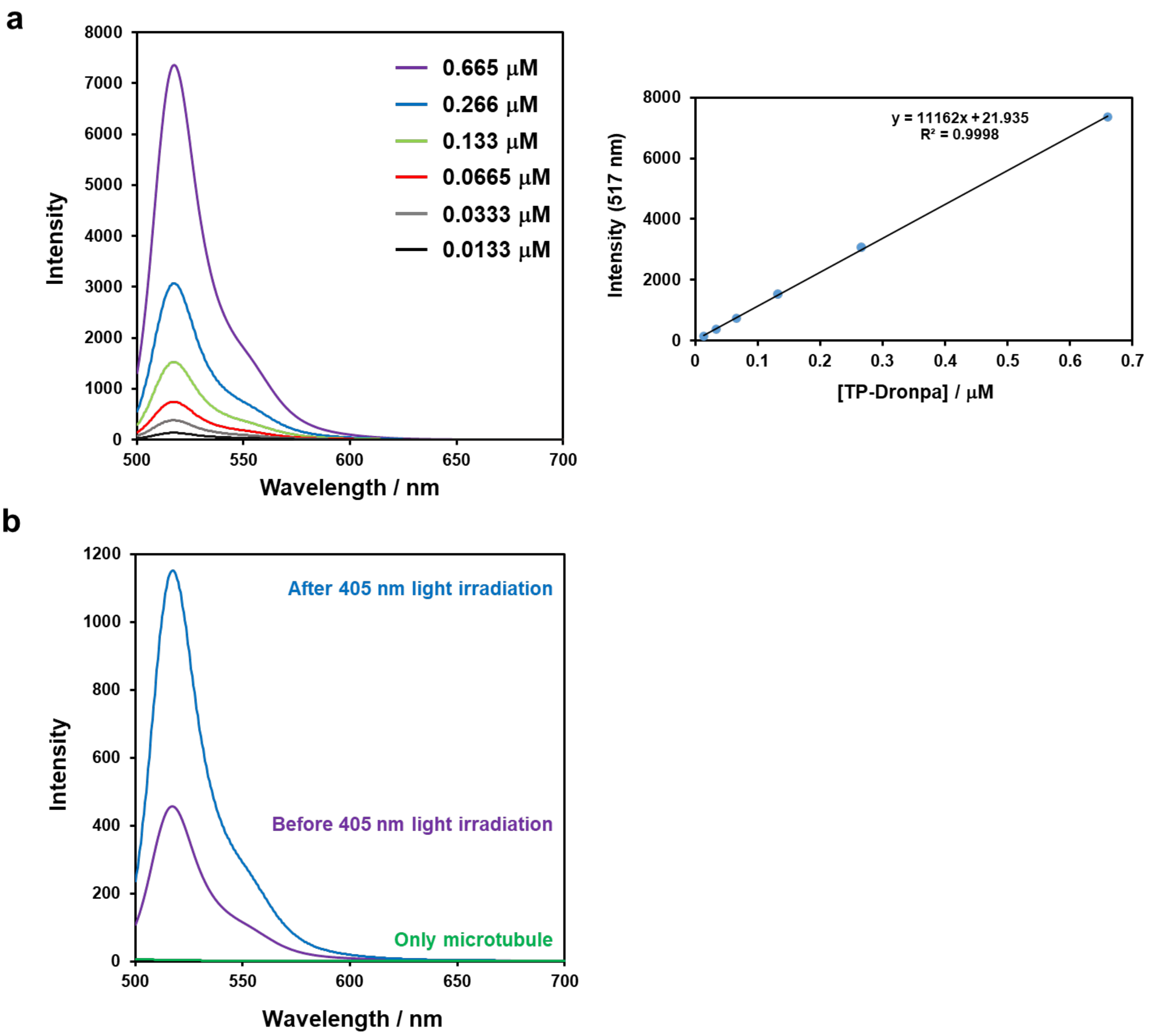


**Figure S9.** Conversion ratio of monomeric TP-Dronpa to tetrameric state on microtubules. (a) Fluorescence spectra of tetrameric TP-Dronpa with different concentration (left) and calibration curve based on the fluorescence intensity (right). (b) Fluorescence spectra of monomeric TP-Dronpa bound with microtubules before 405 nm light irradiation (purple), after 405 nm light irradiation (blue), and only microtubules (green).

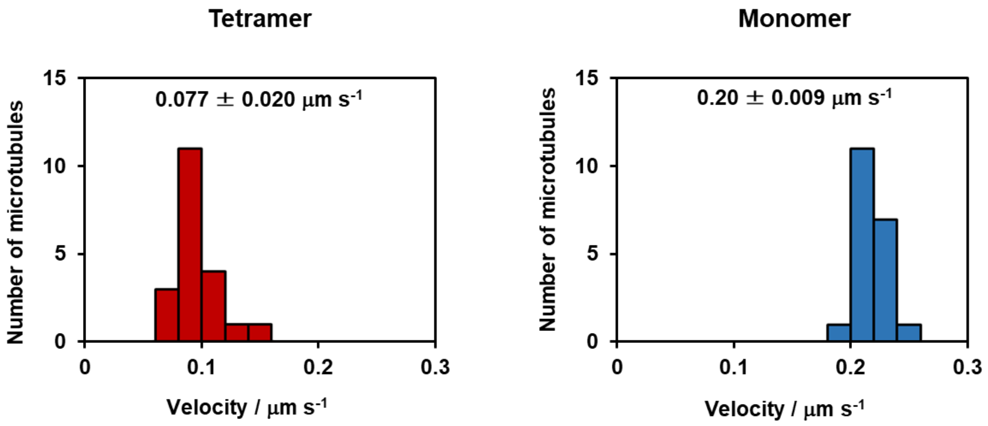


**Figure S10.** Histogram of the velocity of the microtubules with tetrameric or monomeric TP-Dronpa in the presence of methylcellulose ($N = 20$).

**Movie S1.** A movie that illustrates motility of microtubule aster-like structures formed by tetrameric TP-Dronpa (Figure 3b, c). The movie is 100 times faster than the original speed. Scale bar, 30 µm.

**Movie S2.** A movie that illustrates motility of microtubules with monomeric TP-Dronpa. The movie is 100 times faster than the original speed. Scale bar, 30 µm.

**Movie S3.** A movie that illustrates motility of microtubules after conversion of TP-Dronpa bound microtubules from tetrameric to monomeric state by 505 nm light irradiation for 1 h. The movie is 100 times faster than the original speed. Scale bar, 10 µm.

**Movie S4.** A movie that illustrates motility of microtubules after conversion of TP-Dronpa bound microtubules from monomeric to tetrameric state by 405 nm light irradiation for 30 s. The movie is 100 times faster than the original speed. Scale bar, 10 µm.

**Movie S5.** A movie that illustrates motility of microtubules under depletion condition. Tetrameric TP-Dronpa incorporated-microtubules show swarming movement (left). Monomeric TP-Dronpa incorporated-microtubules move individually (right). The movie is 300 times faster than the original speed. Scale bars, 30 µm.

**Movie S6.** A movie that illustrates motility of microtubules with monomeric TP-Dronpa for 5 min, and then the sample was irradiated at 405 nm for 30 s to convert the monomeric form to the tetrameric state. The movie is 300 times faster than the original speed. Scale bars, 30 µm.
